# The influence of sex on the urogenital microbiome of rhesus monkeys

**DOI:** 10.1101/555771

**Authors:** L. K. Hallmaier-Wacker, S. Lueert, C. Roos, S. Knauf

## Abstract

The vaginal microbiome of nonhuman primates (NHPs) differs substantially from humans in terms of *Lactobacillus* abundance, overall taxonomic diversity, and vaginal pH. Given these differences, it remains unclear in what way the NHP genital microbiome protects against pathogens, in particular sexually transmitted diseases. Considering the effect that microbiome variations can have on disease acquisition and outcome, we examined endogenous and exogenous factors that influence the urogenital microbiome of captive rhesus monkeys. The male urethral (n=37) and vaginal (n=194) microbiome of 11 breeding groups were examined in a cross-sectional study. During lactation and menstruation, the vaginal microbiome becomes significantly more diverse and more similar to the microbes observed in the male urethra. Group association and cage-mate (sexual partners) relationships were additionally associated with significant differences in the urogenital microbiome. Our results demonstrate that microbiome considerations are necessary in order to make informed selection of NHPs as translational animal models.

## Introduction

In recent years there has been an increased interest in the microbiome of nonhuman primates (NHPs) for evolutionary, experimental, and conservation purposes. However, microbiome considerations are currently not used to refine and reduce experiments with NHPs, despite increasing evidence that the microbiome in humans can influence disease progression (reviewed by (*1*)). Of the NHPs animal models, the Asian rhesus monkey (*Macaca mulatta*) and long-tailed macaque (*Macaca fascicularis*) are the most extensively utilized species *(2–4)*. In laboratory settings, rhesus monkeys cycle year-round, have a reproductive cycle that is similar to that of humans and experience similar changes in the hormonal levels during sexual cycle, pregnancy and post-partum (*5–7*). Therefore, the vagina of rhesus monkeys has been used to model the human vaginal epithelium and study sexual transmitted infections (STIs) (*8*). For example, rhesus monkeys have been extensively used to study the disease acquisition and outcome of simian-/human immunodeficiency virus (SIV/HIV) (*3, 9*). In a study on SIV susceptibility, estrogen treatment in rhesus monkeys protected female rhesus monkeys from the sexually transmitted infection (*3*). *Smith et al*. propose that not just the thickening of the vaginal epithelium but also a potential change in vaginal microenvironment may have led to the observed effect under the influence of high estrogen levels (*3*).

Many studies have laid the groundwork in characterizing the genital microenvironment of various species of captive and wild NHPs (*2, 10–12*). Unlike the vaginal microbiome of humans, which is often dominated by a single *Lactobacillus* species (*13*), NHPs, including rhesus monkeys, harbor a diverse set of vaginal microbes (*2, 11*). In humans, the acidic nature of the vaginal flora (pH≤4.5) protects women against STIs (*14*). The vaginal microbiome of NHPs on the other hand, has a low abundance of *Lactobacillus* (<2% of microbiome), an overall higher taxonomic diversity, and a near neutral vaginal pH (*2, 11, 12, 15*). Considering these differences, it currently remains unclear in what way the vaginal microenvironment of rhesus monkeys protects against infectious diseases. Additionally, despite increasing evidence that sexual exposures can alter the composition of the human genital flora (*16, 17*), the urethral microbiome of male NHPs remains largely uncharacterized. A better understanding of factors that influence the rhesus monkey genital microbiome of both male and female animals in health and disease is thus warranted.

In this study, we investigate the genital microbiome of a large breeding colony of rhesus monkeys at the German Primate Center. To identify endogenous and exogenous factors that influence the microbiome, we examine the genital microbiome in the breeding colonies in the context of age, breeding group association, social rank, body mass, and long-term health status. We studied both, the vaginal microbiome of female and the urethral microbiome of male rhesus monkeys which has not been done in previous studies. We are therefore able to compare bacterial composition between male and female animals in a single cohort of rhesus monkeys and found that during menstruation and lactations the vaginal microbiome shifts towards the male urethral microbiome.

## Results

We examined the vaginal microbiome of 194 female rhesus monkeys and the urethral microbiome of 37 male rhesus monkeys housed at the breeding facility of the German Primate Center (data file S1). The mean age, number of breeding groups, and other characteristics of the sampled animals are shown in Table 1. The V4 region of the bacterial 16S rRNA gene was amplified and sequenced on the Illumina MiSeq platform, generating a total of 14,571,505 unfiltered reads with a mean read count of 48,630 reads per sample (±15,461 SD) after quality filtering. Sequences were rarefying to 11,371 sequences per sample and clustered into Operational taxonomic units (OTUs) based on the 97% similarity threshold. We first examined the microbiome of the male urethra and vagina separately and then compared composition similarities between male and female animals.

**Table 1:** Characteristics of sampled rhesus monkeys in this study.

|  | Female (n=194) | Male (n=37) |
| --- | --- | --- |
| Mean age, years | 10.0 ± 4.9 | 7.9 ± 5.6 |
| n Geriatric (>19 years) | 10 | 1 |
| n Adult (5-19 years) | 168 | 17 |
| n Juvenile (< 5 years) | 16 | 19 |
| Breeding groups | 11 | 10 |
| Vaginal pH, mean ± SD | 6.4 ± 0.7 (n=138) | N/A |
| % EVC* | 76.3% (n=148) | N/A |
| n Phase 1 | 44<sup>†</sup> | N/A |
| n Phase 2 | 57<sup>†</sup> | N/A |
| n Phase 3 | 47<sup>†</sup> | N/A |
| % Lactating | 30.4 (n=59) | N/A |
| % Dominant male | N/A | 40.5 (n=15) |

\*Cytology phases classification described in Detail in Methods and Supplementary Table S2;
<sup>†</sup> For subdivision of sexual cycle phases by lactating status see Table S2

Appropriate control samples and a mock community (Microbial mock community, HM-280, Biodefense and Emerging Infectious Research (BEI) Resources, Manassas, USA) were included in the sequencing run. Using the mock community, the observed error rate for the run was found to be 0.036%. The collected control samples showed that contamination was highest during sample collection procedures, while controls taken during amplification procedures in the laboratory yielded only minimal read counts (Fig. S1A). Taxa plots of control samples that were taken during sampling at two different breeding units, show that the relative abundance of contaminant OTUs was similar between the units (Fig. S1B).

### The vaginal microbiome is significantly altered during lactation and menstruation

We investigated the vaginal microbiome (n=194), vaginal pH (n=138), and sexual cycle phase (n=148) of clinically-healthy, reproductively-active rhesus monkeys housed in eleven breeding groups (Table 1). None of the females showed signs of pregnancy, as defined by transabdominal palpation. At the time of sampling 30.4% of the animals were lactating. In addition to lactation, we characterized the sexual cycle phase using exfoliative vaginal cytology (EVC) (see Methods, Table S1). The sexual cycle phase (P1-P3) of non-lactating females were evenly distributed with 35.6% in an ovulatory phase (P1), 41.3% in an intermediate phase (P2) and 23.1% in a menstruation-like phase (P3) (Table S1). For lactating females, 52.3% of the animals were in a menstruation-like phase, 31.8% in an intermediate phase, and 15.9% in the ovulatory phase (Table S1). For the purpose of the microbiome analysis, lactation status and sexual cycle phases were analyzed independently.

Overall, a mean of 219.8±160.7 (unless otherwise stated all values are given in mean ± SD) OTUs were observed in the vaginal microbiota of the rhesus monkeys. The most abundant genus was *Prevotella* with a mean abundance of 20.5±16.4%. Different OTUs were identified as *Prevotella*, indicating that a diverse set of species from this genus were present (Fig. 1). *Porphyromonas (*9.5±9.9%), *Streptobacillus* (9.1±13.4%) and an unclassified genus of the family *Ruminococcaceae* (9.5±7.3) were the other dominant taxa in the otherwise diverse community (Fig. 1).

**Fig. 1:**
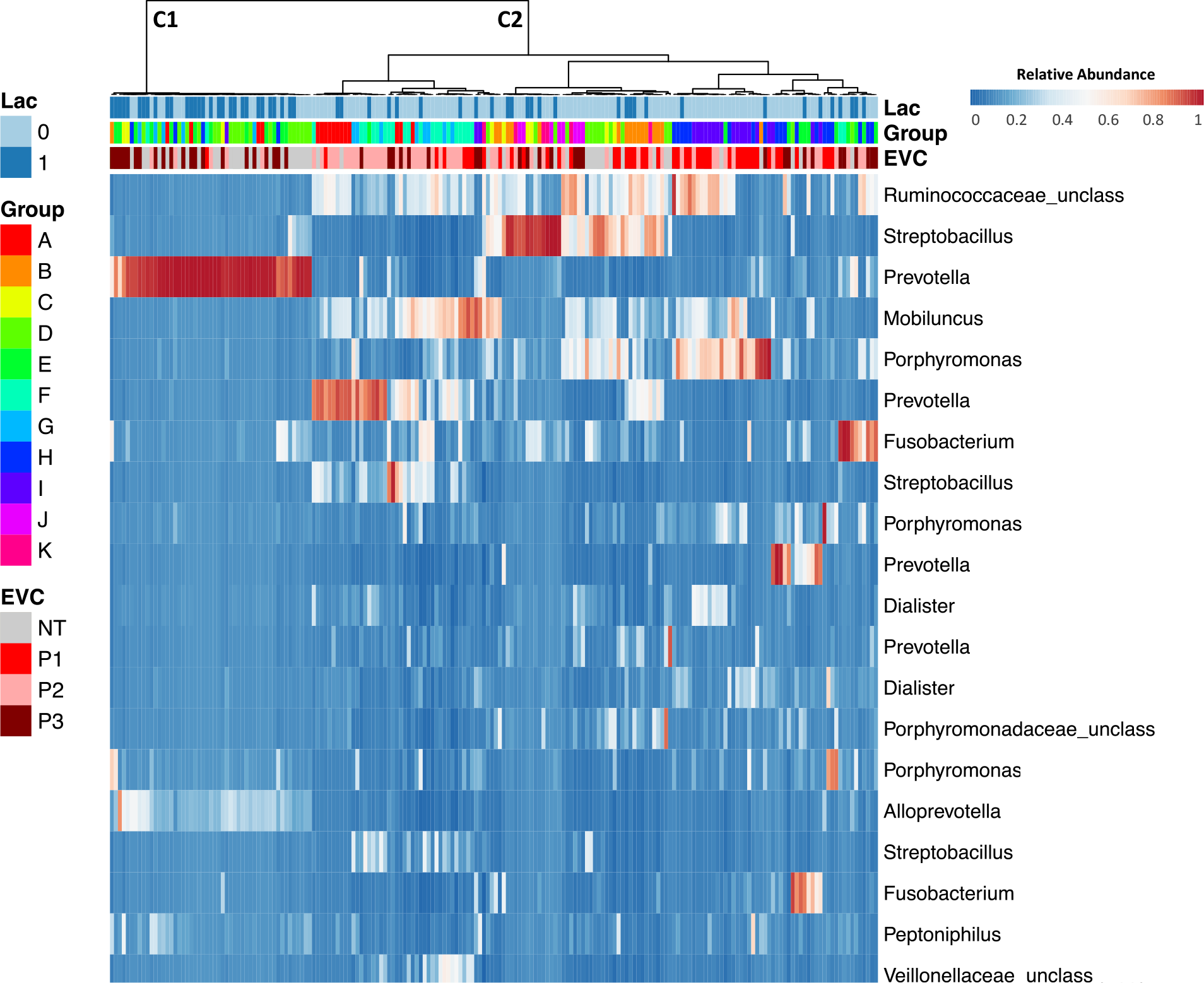
Heatmap of the relative abundance of microbial taxa identified in the vaginal microbiota of rhesus monkeys in multiple breeding groups. Ward linkage clustering of samples based on the composition and relative abundance of the 20 most abundant OTUs in the vaginal microbiota. Genus-level bacterial classification of OTUs shown with the percent of sequences that classified with the specific genus. Lactation status (1: lactating and 2: non-lactating), group association (A-K) and EVC (sexual cycle phases) (P1: ovulatory phase, P2: intermediate stage, P3: menstruation-like and NT: not tested) of each sample are shown beside the heatmap. C1 and C2 indicate two main clusters in the ward linkage clustering. See Table 1 for sample size composition.

Lactation status and sexual cycle phase strongly correlated with the OTU richness (identified absolute number of taxa) and evenness (inverse Simpson index; Fig. 2). Lactating females had a significantly higher OTU richness (<p=0.0001 [Mann-Whitney t-test]) and the bacterial taxa were significantly more evenly distributed (p=0.0001 [Mann-Whitney t-test] than in non-lactating females (Fig. 2A-B). Similarly, animals in a menstruation-like (Phase 3, Table S1) sexual cycle phase had a significantly higher OTU richness (<p=0.0001 [Kruskal-Wallis test]) and were significantly more evenly distributed (p=0.001 [Kruskal-Wallis test]) than animals in the ovulatory (Phase 1) or intermediate phase (Phase 2; Fig. 2C-D). A heatmap of the relative abundance of the 20 most common OTUs shows that lactating animals and animals in menstruation-like sexual phase clustered separately from other animals (Fig. 1). Vaginal bacterial communities from these animals clustered prominently in cluster 1 (C1) and are characterized by different bacterial taxa than the cluster 2 (C2; Fig. 1). Of the ten most abundant OTUs, *Provella*, *Mobiluncus, Porphyromonas* and an unclassified genus of the family *Ruminococcaceae* were significantly different in the lactating and menstruation-like animals (Fig. S2A-B). These cluster differences were confirmed by significant differences in the unweighted UniFrac distances, which are visualized on the principal coordinates plot along axis 1 (23.1%) (Fig. 3A-B). Pairwise AMOVA confirmed that the differential clustering of lactating and menstruating-like animals resulted in significantly different community structures (p=0.001).

**Fig. 2:**
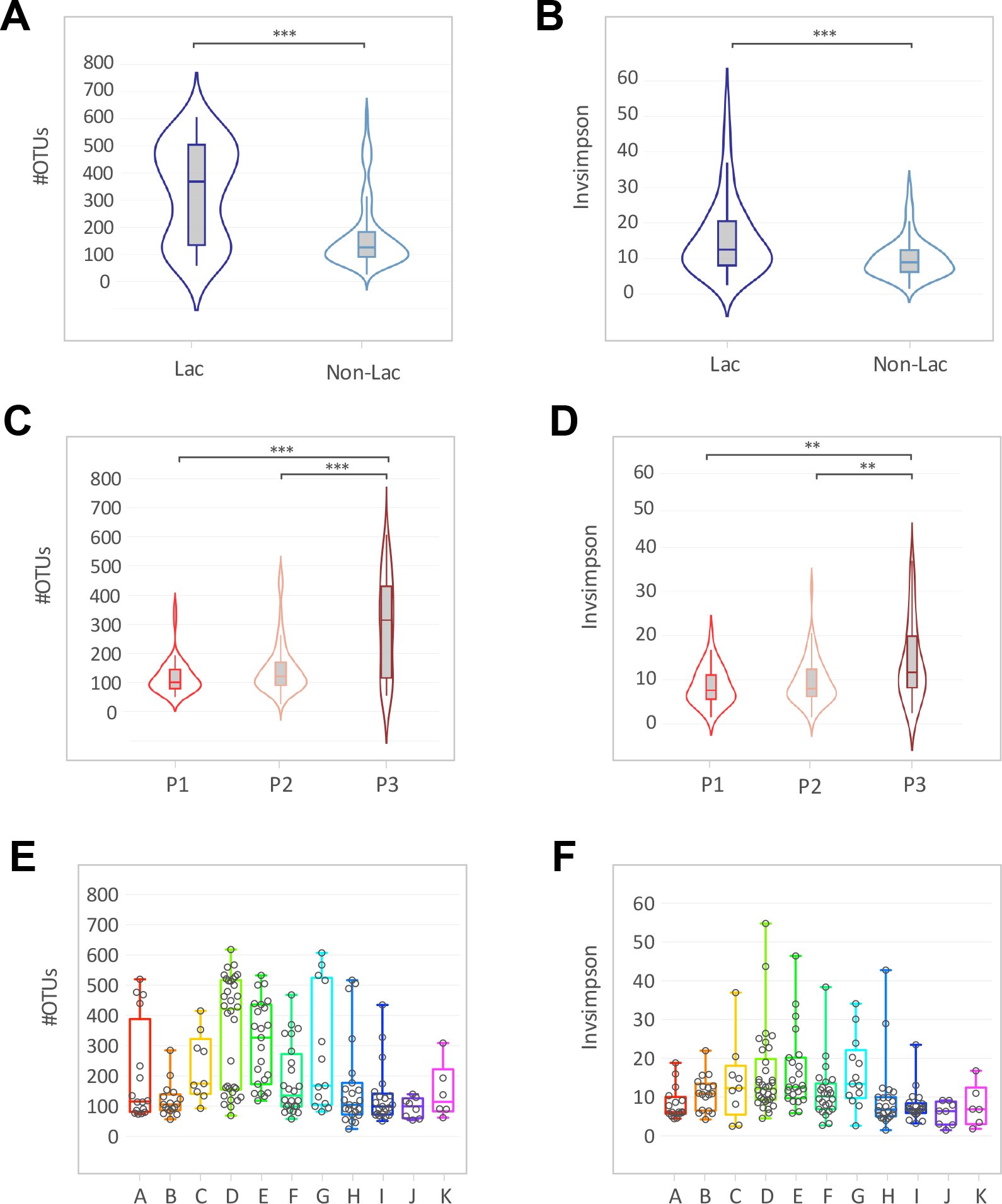
Alpha diversity measurements for vaginal microbiome of female rhesus monkeys. Violin plots of the observed OTUs and InvSimpson index clustered based on (**A/B**) lactation status (Mann-Witney t-test, ***p≤0.0001) and (**C/D**) sexual cycle phases (P1: ovulatory phase, P2: intermediate stage, P3: menstruation-like) (Kruskal-Wallis test, **p≤0.001, ***p≤0.0001). (**E/F**) Boxplots (median ± range) of the observed OTUs and InvSimpson index clustered of breeding groups (groups association: A-K). See Table 1 for sample size composition.

**Fig. 3:**
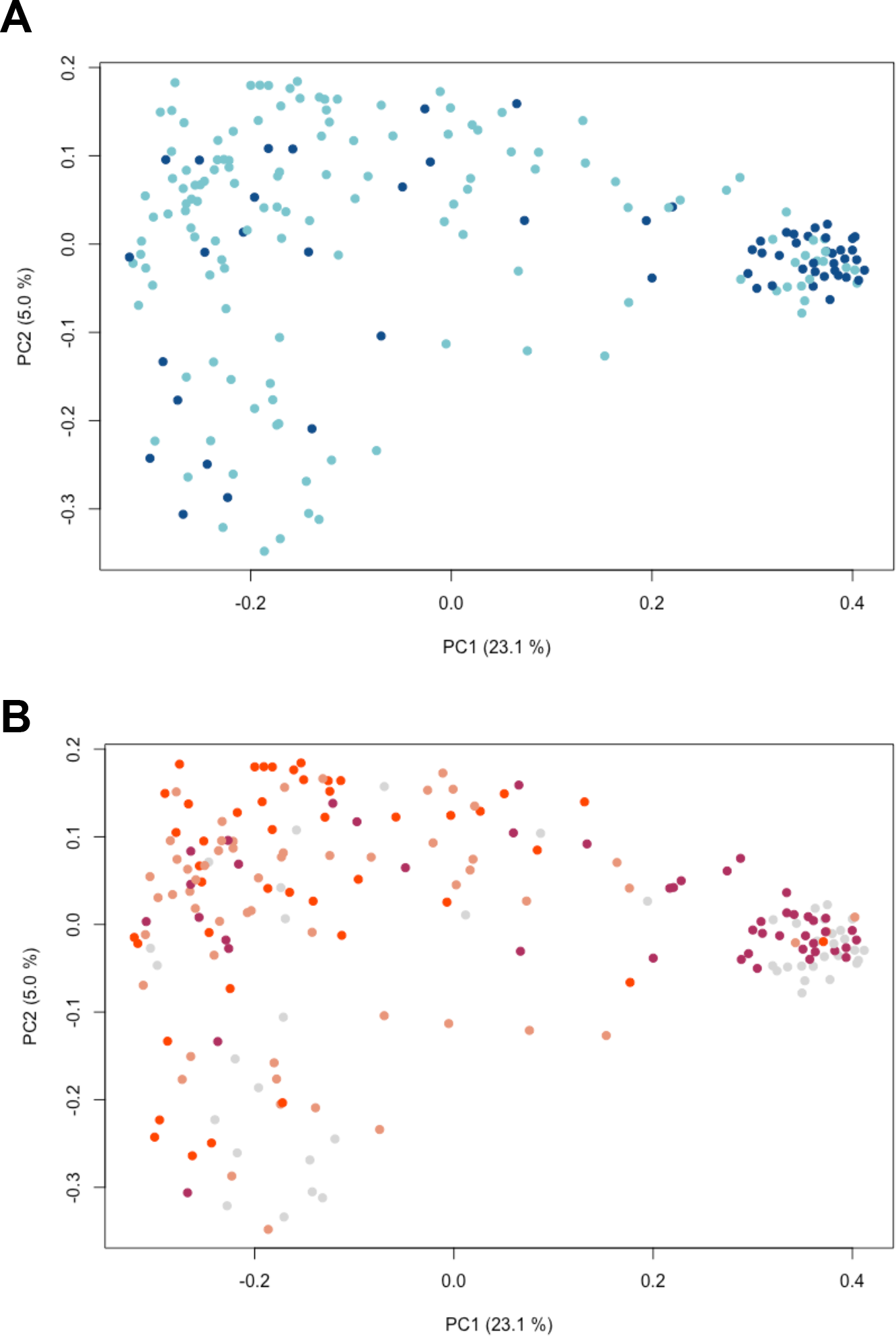
The vaginal microbiome of menstruating-like and lactating females clusters separately. Principal coordinates analysis (PCoA) of vaginal samples colored by (**A**) lactation status (lactating (dark blue) and non-lactating (light blue) and (**B**) sexual cycle status (P1: ovulatory phase (red), P2: intermediate stage (pink), P3: menstruation-like (dark red) and NT: not tested (gray)). Distances between samples were calculated using the unweighted UniFrac metrics. See Table 1 for sample size composition. Fig. S5 shows the corresponding PCoA plot classified by group association and age classification.

In order to examine an additional functional variable of the vaginal microbiota, we tested the vaginal pH at the time of sampling using pH-indicator paper. The mean overall vaginal pH of the sampled animals was found to be 6.4±0.7 (Table 1). The vaginal pH of lactating females was significantly higher than that of non-lactating females (>p=0.0001 [Mann-Whitney t-test], Fig. S3A). Similarly, animals in menstruation-like sexual phase had a higher vaginal pH compared to individuals in the other sexual cycle phase (>p=0.0001 [Kruskal-Wallis test], Fig. S3B).

### Breeding groups influence the vaginal microbiome

Aside from lactation status and cycle phase, we examined if the vaginal microbiome is shaped by age and breeding group. Animals were subdivided into juveniles (<5 years), adults (5-19 years), and geriatric (>19 years). Age was not found to have an influence on alpha diversity (Fig. S4) and beta diversity (p=0.155 [AMOVA; Fig. S5A). The group association, in contrast, significantly correlate with the OTU richness (>p=0.0001 [Kruskal-Wallis test], Fig. 2E) and evenness (>p=0.0001 [Kruskal-Wallis test], Fig. 2F). The breeding group association of each sample can be seen on the heatmap of the 20 most abundant OTUs (Fig. 1). Additionally, we observed a significant difference in unweighted UniFrac distances when considering all breeding groups (<p=0.001 [AMOVA; Fig. S5B). Pairwise comparisons of alpha and beta diversity measurements between individual breeding groups was, however, not significant for all tested groups.

### Influence of social rank in adult male rhesus monkeys

We characterized the urethral microbiome of clinically-healthy, reproducing male rhesus monkeys housed in ten different breeding groups (Table 1). Overall, the urethral microbiome of male rhesus monkeys is composed of a diverse community of microbes, with a mean of 481.3±127.0 OTUs observed. On a phylum level, *Firmicutes* (54.1±8.3%), *Bacteroidetes* (25.5±9.3%), *Proteobacteria* (9.0±6.7%) and *Actinobacteria* (6.6 ± 3.2%) made up 95% of the identified sequences. The four dominant phyla were present in all 37 samples. On the genus level, the bacterial community is diverse with no single dominating OTU (Fig. 4). The most abundant genus in the male rhesus monkey urethra was *Prevotella* with a mean abundance of 14.4±9.7% followed by *Porphyromona* (7.5±6.6%) and *Ezakiella* (7.3±6.8%). We examined if a dominance rank in the breeding group shaped the urethral microbiome. The highest-ranking male was classified as the dominant male. Each breeding group had one dominant male with the exception of group H (n=4) and I (n=3), which were further divided into subgroups within a single housing unit. Dominant males neither differed from other males in the OTU richness (p=0.145, [Mann-Whitney test], Fig. 4A) nor the evenness (p=0.453 [Mann-Whitney test], Fig. 4B). Pairwise AMOVA of unweighted UniFrac distances found that dominance rank had no effect on community structure (p=0.123). A dendrogram of unweighted UniFrac distances shows that dominant animals did not cluster separately from other animals (parsimony analysis, p=0.768; Fig. 4C). OTU richness and evenness measurements of each breeding group are shown in Fig. S6. We note here, that the sample size of male animals in each breeding group were low (n=1 to 4 animals). Therefore, statistical analysis was not performed to examine breeding group differences.

**Fig. 4:**
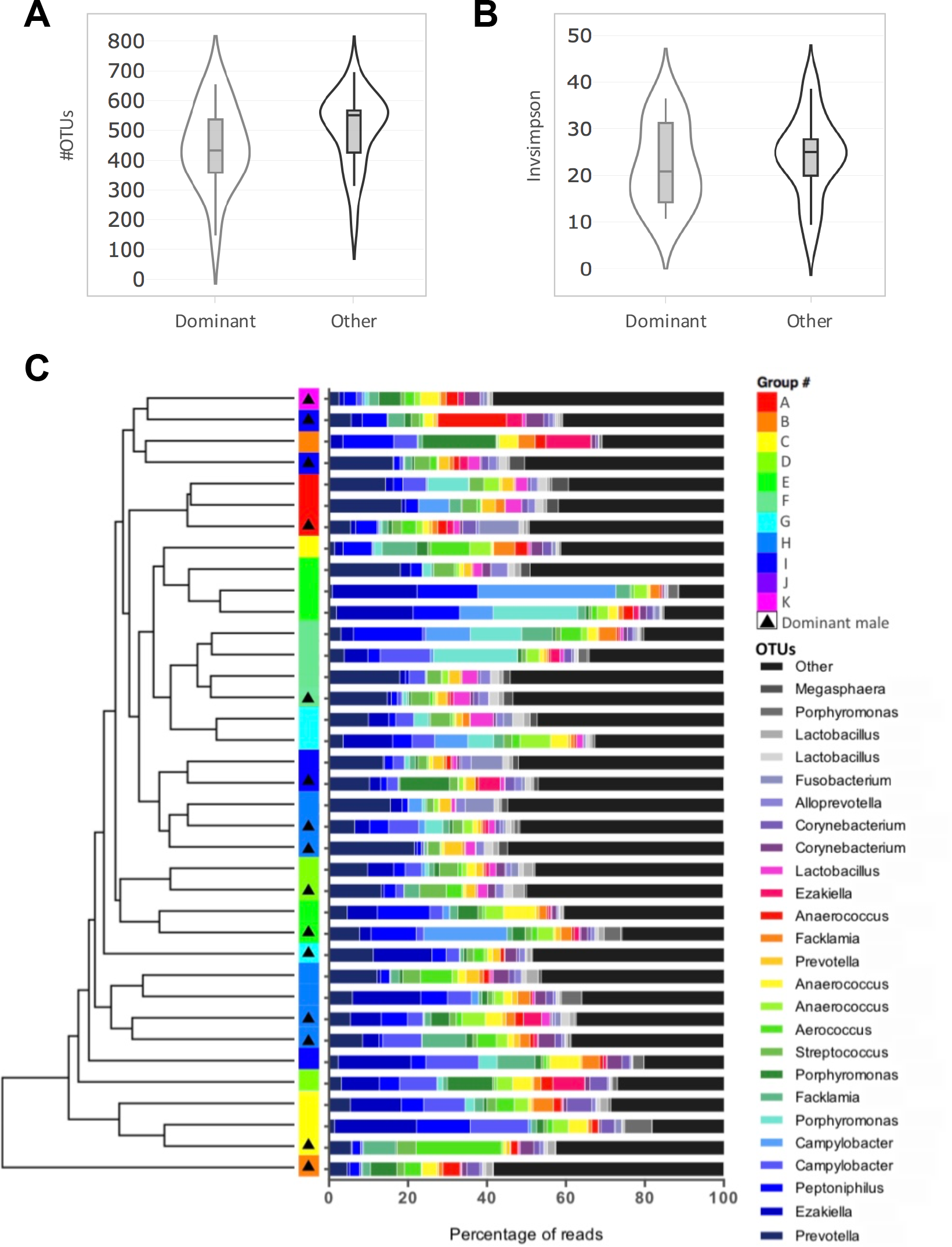
The male urethral microbiome of dominant and non-dominant (other) rhesus monkeys does not differ significantly. Violin plot of (**A**) the number of OTUs and (**B**) the inverse Simpson index for dominant and non-dominant (other) males (Mann-Witney t-test). (**C**) UPGMA clustering on unweighted UniFrac including taxa plots showing the relative abundance of the 25 most abundant OTUs in percentage of reads. Genus-level bacterial classification of OTUs shown legend with the percent of sequences that classified with each genus. Group and dominance rank are shown accordingly. See Table 1 for sample size composition.

### Lactating and menstruation female have a more similar microbiome to the male urethra

On the phylum level, *Firmicutes* and *Bacteroidetes* dominated both microbiomes, making up 80.1±6.9% in the male urethra and 69.9±17.8% in the vagina. Yet, *Fusobacteria*, the third most abundant phylum in the vagina (14.4±16.1%), only made up 1.9±3.3% in the male urethra. The most abundant genus across the dataset for both, male and female genital microbiome was *Prevotella*. Several OTUs cluster into this bacterial genus and the most abundant *Prevotella* OTUs (mean abundance of 6.0±8.7%) was found in 226 out of 231 animals. Other *Prevotella* OTUs were less abundant and only dominant in some samples (Fig. 1: vaginal microbiome and Fig. 4: male urethral microbiome).

To further examine similarities between the vagina and male urethra, overall OTU richness and evenness was compared. As we previously observed a significant difference in alpha and beta diversity of the vaginal microbiome based on lactation status and sexual cycle phase, these variables were plotted separately (Fig. 5). The male urethra had a significantly higher OTU abundance compared to non-lactating and non-menstruation-like (ovulatory and intermediate phase) animals (<p=0.0001 [Kruskal-Wallis test], Fig. 5A/B). Contrary, menstruation-like (P3) and lactating female rhesus monkeys showed no significant difference in the number of OTUs compared to the male urethra microbiome (>p=0.05 [Kruskal-Wallis test], Fig. 5A/B). Similarly, inverse Simpson index measurements were significantly different between males and non-lactating and non-menstruation-like (ovulatory (P1) and intermediate phase (P2)) animals (<p=0.0001 [Kruskal-Wallis test], Fig. 5C/D). Inverse Simpson index measurements were not significantly different between males and lactating and menstruation-like animals (>p=0.05 [Kruskal-Wallis test], Fig. 5C/D). To examine, if the trend in the alpha diversity could be observed in the overall bacterial composition, pairwise unweighted UniFrac distances were calculated between the male urethral microbiome and the vaginal microbiome. More similar microbiomes resulted in a smaller calculated UniFrac distances and vise versa. The UniFrac distances were grouped in violin plots based on either lactation status (Fig. 5E) or sexual cycle phase (Fig. 5F). We found that the bacterial composition of the vaginal microbiome of lactating and menstruating-like animals (P3) was significantly more similar to that of the male urethra microbiome (Fig. 5E/F).

**Fig. 5:**
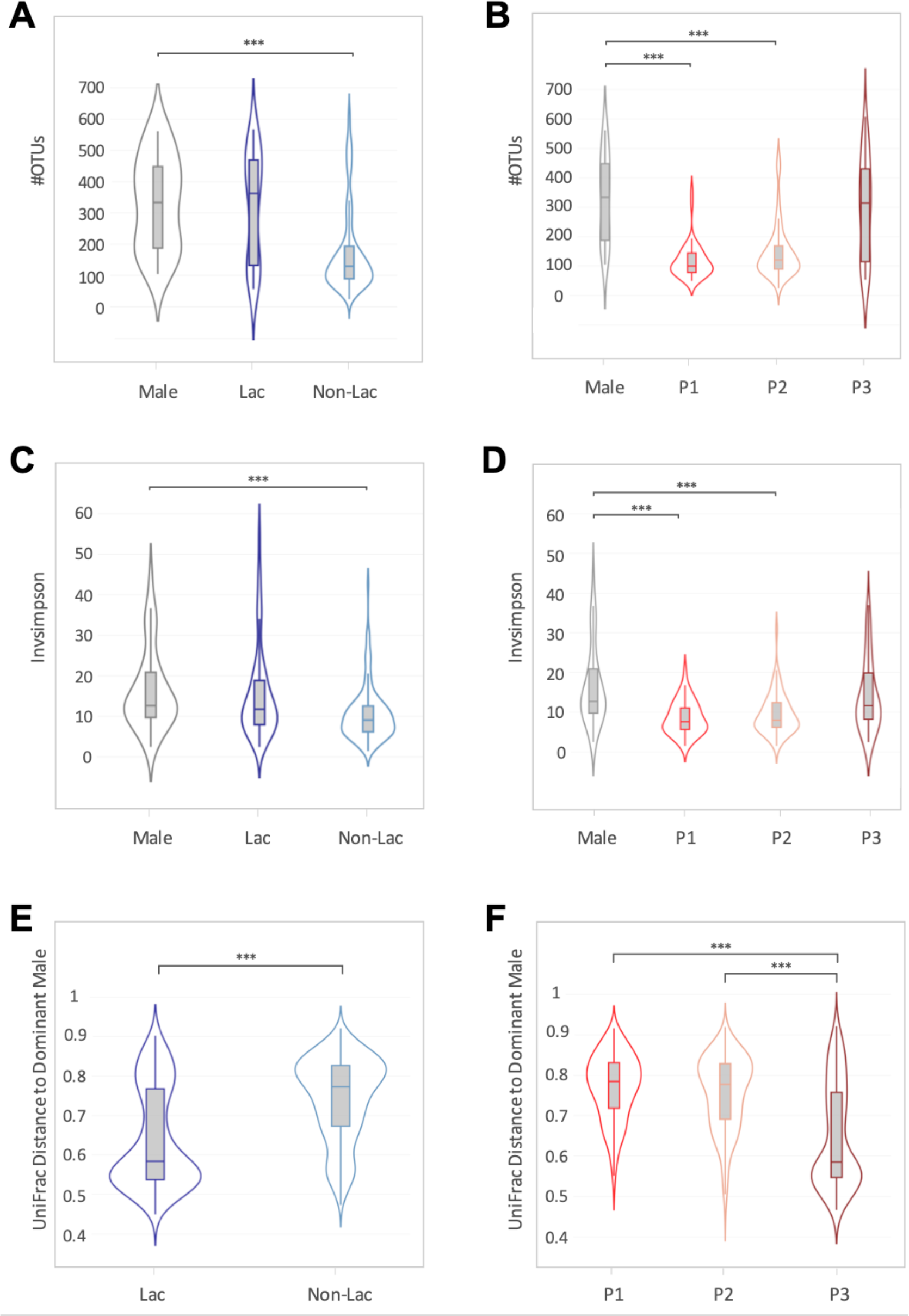
The vaginal microbiome of lactating and menstruating females is more similar to the urethral microbiome. Violin plot of the number of OTUs (**A**) in the male urethra and vaginal microbiome of lactating and non-lactating females and (**B**) in the male urethra and vaginal microbiome of females in three sexual cycle phases (P1: ovulatory phase, P2: intermediate stage, P3: menstruation-like) (Kruskal-Wallis test, ***p≤0.0001). Violin plot of the inverse Simpson index for (**C**) male urethra and vaginal microbiome of lactating and non-lactating females and (**D**) male urethra and vaginal microbiome of females in three sexual cycle phases (Kruskal-Wallis test, ***p≤0.0001). (**E/F**) Violin representations showing unweight UniFrac Distance of each female to the dominant male in each group. Data is plotted by (**E**) lactation status (Kruskal-Wallis test, ***p≤0.0001) and (**F**) sexual cycle phase (Mann-Witney t-test, ***p≤0.0001). See Table 1 for sample size composition.

### Cage-mates are more similar in their genital microbiome

In order to assess if sexual contact shapes the genital microbiome of the captive rhesus monkeys, pairwise unweighted UniFrac distances were calculated between the male urethral microbiome of the dominant male in each group and the vaginal microbiome. Females and males of the same breeding group were considered cage-mates and thus potential sexual partners. UniFrac distances were grouped into violin plots as ‘cage-mates’ or from ‘other breeding groups’ (no sexual contact possible) (Fig. S7A). Cage-mates were found to be significantly more similar in the bacterial composition compared to non-cage-mates (<p=0.0001 [Mann-Whitney test], Fig. 7A). As we observed a significant difference in lactation status and sexual cycle phase, these variables were additionally plotted in separate paired-violin plots to examine cage-mate differences for each group (Fig. S7B/C). Cage-mates were found to be significantly more similar in the bacterial composition for lactation, menstruation-like and ovulatory phase animals (Fig. S7B/C). Cage-mate similarity was not observed for the non-lactating group or for animals in an intermediate sexual phase (P2; Fig. S7B/C).

## Discussion

Considering the effect that microbiome variation can have on disease acquisition and outcome (*1*), we examined endogenous and exogenous factors that influence the urogenital microbiome of captive rhesus monkeys. The inclusion of appropriate controls (Fig. S1) and the large sample size confer confidence in our study. However, based on our cross-sectional study design we were limited in drawing causal relationships between factors and variations in the genital microbiome. Nevertheless, our results urge for the inclusion of microbiome analysis in the selection and experimental use of rhesus monkeys as indicated by the differences between the vaginal microbiome during lactation and sexual cycles phases.

We showed that during the endocrine state, which causes the female to lactate and menstruate the bacterial composition shifts towards a more diverse community (Fig. 1-3). As reported previously, we confirmed that the mean vaginal pH of rhesus monkeys (6.4±0.7) is significantly higher than that found in humans (*2*) (Fig. S3). Instead of the *Lactobacillus-* dominance observed in women (reviewed by (*18*)), the vaginal microbiome of captive and wild NHPs harbor a more diverse set of bacteria (Fig. 1) (*2, 11, 12, 15*). Our study shows that in captive rhesus monkeys the already diverse bacterial community shifts to an even more diverse and significantly different bacterial composition during lactation and the menstruation-like phase (Fig. 2-3). There has been some discussion if the sexual cycle influences the vaginal microbiome of captive NHPs. While a previous study on captive baboons (*Papio anubis*) found no difference in the vaginal microbiome of cycling females (*15*), a recent study in wild baboons (*Papio cynocephalus*) reported that the ovarian cycle phase and the reproductive state shaped the vaginal microbiome (*12*). Both of these studies used visual assessment of perivulvar swellings to determine the sexual cycle phase (*12, 15*). Inconsistent classification of these phases in the two studies in combination with a low sample sizes may explain the difference in the outcome of both studies. Instead of using perivulvar swellings, we performed vaginal exfoliative cytology to classify the animals into three sexual cycle phases ((*19, 20*); Table S1)). Vaginal exfoliative cytology reflects the current state of the vaginal epithelium and therefore serves as a reliable marker for the sexual cycle phase (*19, 20*). The even distribution of all three sexual cycle phases in non-lactating rhesus monkeys (Table S1) is indicative of a healthy reproductive community. Using cytology as a marker of sexual cycle phase, this study supports *Miller et al*.’s finding that ovarian cycle phase (menstruation-like) and reproductive state (lactation) shifts the vaginal microbiome in NHPs (*12*). Similar changes have been reported in temporal and cross-section studies in women during menstruation and post-partum (*13, 21*), where it has been demonstrated that the vaginal microbiome shifted from a *Lactobacillus*-dominant state towards a more diverse bacterial composition (*13, 21*). Despite the remarkable differences in bacterial species composition of the rhesus monkeys and human vaginal microbiome, it is interesting that similar factors (e.g. hormonal changes) seem to influence the vaginal microbiome. This is supported by our finding that the observed changes in bacterial vaginal diversity in the rhesus monkeys coincide with changes in the pH, a functional measurement of the vaginal ecosystem. Understanding the causal relationships that regulate and control changes in vaginal ecosystem is a current challenge for both NHP and human microbiome studies (reviewed by (*22*)) and warrants further investigation.

It has been proposed that hormonal fluctuations during the sexual cycle, pregnancy and post-partum shape the vaginal microbiome (reviewed by (*18*)). Both lactation and menstruation are marked by hormonal changes in the vagina, which may be indirect or direct driving factors for the shift in vaginal microbiome observed in this study (Fig. 1-3). Studies on SIV susceptibility in rhesus monkeys have shown that hormone treatment can lead to an altered susceptibility (*3, 9*). During high levels of estrogen, changes in the vaginal epithelium, including changes in vaginal microenvironment, may have a protective effect (*3*). The less diverse vaginal microbiome and lower overall pH (Fig. S3) found in this study during ovulatory phase supports an important role of the vaginal microenvironment in infectious disease acquisition. Further investigations are necessary to examine the causal relationship between hormone levels, changes in the NHP vaginal microbiome, and susceptibility to pathogens. However, it has become clear, that a more holistic understanding on host-pathogens interactions is required for the interpretation of animal experiments as host factors can influence the microbiome and vice versa (reviewed by (*1*)).

We examined the male urethral microbiome of the rhesus monkeys to further compare the genital microbiome of females and males in a single breeding unit. To our knowledge, there has been no studies on the urethral microbiome of wild or captive NHPs to date. Four bacterial phyla, *Firmicutes, Bacteroidetes*, *Proteobacteria*, and *Actinobacteria*, compose the majority of identified sequences in the urethra. On the phylum level, the urethral microbiome of the male rhesus monkeys were similar to that reported in humans with *Firmicutes* making up the largest proportion (*23*). In our male animals, notable urethral taxa include *Prevotella*, *Porphyromona*, and *Ezakiella*, have all been previously associated with the urinary tract microbiome of adult men (*23–25*). *Prevotella* has been previously detected in the genital microbiome of healthy female rhesus monkeys indicating that this genus plays a residential role in the rhesus monkeys genital microbiome (*10*). In humans, some species of *Prevotella* have been associated with disease states (e.g., bacterial vaginosis (*26*)) while other species can be found in clinically healthy women (e.g., post-partum, (*21*)). Identifying the role of *Prevotella* in NHPs may assist in a better functional understanding of the genital ecosystem. For the urethral microbiome, it is difficult to compare the prevalence of *Prevotella* in the male rhesus monkey to other studies, as there is currently no consensus on the core urethral microbiome, even in humans (*25*). As a result, large scale investigations need to be performed to study the male urogenital microbiome including factors that influence this unique ecosystem in health and disease (*25*).

It has been hypothesized that sexual exposures can alter the composition of the genital flora (*13, 16, 25*). A recent study on sexual partners with bacterial vaginosis (BV), showed that women with BV were significantly more similar to the urogenital microbiome of their partner (*17*). To test if sexual contact affected the genital microbiome of NHPs, we first examined if dominance rank in captive rhesus monkeys shaped the male urethral microbiome. Breeding groups in this study contained a single breeding male, who monopolized the cage-mates in estrus. We did not find that dominance rank shaped the male urethral microbiome (Fig. 4). This may be due to the fact that non-dominant juvenile rhesus monkeys already engage in socio-sexual mounting as a form of play (*27*). Contrary to our findings, sexual history in healthy adolescent men has been reported to be a possible determinant of the urogenital microbiota (*16*). Known sexually transmitted bacteria and taxa associated with the urethral tract of adult men (*23*), were observed rarely in adolescent men (*16*). To further study the effect of sexual contact, we examined the similarity of the genital microbiome in cage-mates (alpha male to females in the same breeding group). We found that overall, cage-mates were significantly more similar to each other compared to non-cage-mates (Fig. S7A). When subdividing cage-mate by lactation status and sexual cycle, the observed cage-mate effect was not seen for non-lactation and intermediate sexual phase animals (Fig. S7B-C). This may be due to an inappropriate subsampling of these two groups. For example, the intermediate sexual phase classification used in the EVC may represent both, proliferative phase and secretory phase, and is therefore an oversimplification. This highlights the limitation of this cross-sectional study in assessing cage-mate similarities. A controlled temporal study is necessary to examine the effect of sexual contract in NHP breeding groups. NHPs can be an advantages model to further examine microbiome similarities in sexual partners (in health and disease) as sexual contact is easily observed and controlled.

A surprising finding of our study was that independent of breeding group association, the bacterial composition of lactating monkeys and/or those in the menstruation-like sexual phase were more similar to the microbiome of the male urethra (Fig. 5). As females in a menstruation-like sexual phase are less attractive to male rhesus monkeys, we presume that the similarity is not caused by recent sexual contact. A possible explanation for this finding is that the altered hormonal state allows otherwise more-suppressed bacteria to dominate the microflora. To understand the cause of the vaginal microbiome shift towards the male urethra microbiome, controlled temporal experiments in NHP would be necessary. Interestingly, a temporal study in humans has shown that the vaginal microbiome post-partum shifts towards the gut microbiome (*21*). The study was able to show that the shift towards the gut microbiome persisted for multiple months and was independent of delivery method (vaginal vs. caesarean). These findings support the notion that during changes in the genital ecosystem (e.g., shifts in hormones), the vagina is more susceptible to ‘foreign’ bacteria. This potentially altered susceptibility should be carefully considered when performing vaginal inoculations in NHPs for future experiments (e.g., HIV).

We found that breeding groups can have an effect on the vaginal microbiome (Fig. 2). Breeding group similarities could be influenced by various factors including host genetics (*28*), differences in group size or cage effects (*29*). Many of these factors could not be properly examined in this study and require planned and controlled animal experiments. In mice, it has been shown that animals kept in the same cage become more similar in microbiome composition over time (*29*). This effect could be studied in captive NHPs by examining microbiome changes in various ecological niches (genital, skin, fecal) during cage transfers. A cage effect in NHPs could have major implications for the use of NHPs as translational animal models. A better understanding of the NHP microbiome could therefore refine animal selection for animal experiments where a higher standardization can lead to reduced animal numbers (*3, 9*). The inclusion of appropriate controls in microbiome studies cannot be stressed enough (*30*). Especially low abundance microbiomes like the urethral microbiome are vulnerable to contaminations during sampling and laboratory analysis (*31*). The inclusion of blank control samples, especially at the site of sample collection, is essential and should be understood as Good Laboratory and Scientific Practice (Fig. S1). Only well-planned and controlled microbiome studies on NHPs are capable of effectively reduce and refine NHP numbers in translational animal models and providing a better understanding of factors that influence microbiomes of NHPs.

## Methods

### Ethical statement

All samples included into this study were obtained from clinically healthy rhesus monkeys that underwent the mandatory annual health check at the German Primate Center between June 2016 and May 2017. Animals were not purposely immobilized to collect samples for this study. Swabs were taken as part of a routine annual health monitoring and tuberculosis screening. Animal were short-term immobilized by trained veterinarians who checked and documented the general health condition of each individual. Sampling included the collection of blood, oral and genital swab samples. The use of the samples was reviewed and approved by the animal welfare and ethics committee of the German Primate Center (EC No. 1-16). All work steps involving the handling of live animals followed the rules of ‘Good Veterinary Practice’.

### Study Design and Animals

Urethral swabs of 37 male and vaginal swabs of 194 female rhesus monkeys were collected. A cross-sectional study design was applied. Samples from apparently pregnant individuals, clinically diseased animals, or animals that received medical treatment within the last 6 months were excluded from analysis. Moreover, we excluded samples from animals below the age of three. Data file S1 provides a detailed overview on the samples analyzed in this study as well as the respective NCBI Sequence Read Archive numbers. Lot numbers for consumables were kept consistent and are reported in the Supplementary Material (Table S2).

### Swab Sample Collection

Immobilized female rhesus monkeys were placed in dorsal recumbency and the area around the vulva was cleaned using 70% ethanol. To facilitate sampling, a sterile silicon tube, 15 mm diameter and 40 mm length, was used to avoid swab contamination with skin or fecal material. A flocked swab (FLOQSwabs, Copan Improve Diagnostics) was moistened using a single drop of sterile physiological saline solution (WDT eG) and was subsequently inserted midway into the vaginal canal. Subsequently the swab was rotated 20-times on the dorsal wall before it was gently removed and transferred into 500 µl of custom-made lysis buffer (10mM Tris, pH 8.0, 0.1M EDTA, pH 8.0 and 0.5% SDS). Samples were kept on ice until transported to the inhouse laboratory facilities where they were stored at −80°C (*30*).

An additional swab was collected to perform an EVC. Briefly, the swab was rolled onto a microscope glass slide after which it was allowed to air-dry. Slides were then stained with a Romanowski stain (Diff-Quik) and subsequently examined under the microscope by two independent investigators (*19*). Cytological scoring was performed as previously described by *McLennan et al*. (*20*). The maturation index was calculated by counting 100 representative epithelial cells, which were scored according to their cell type. Briefly, parabasal cells were assigned a value of 0, intermediate cells a value of 0.5, and superficial cells a value of 1. Based on the cumulative maturation score, the animals were categorized into three stages (ovulatory phase (P1), intermediate phase (P2), and menstruation-like phase (P3); see Supplementary Table 2).

The vaginal pH was measured using a swab which was inserted midway into the vagina and then rolled onto a pH-indicator paper (Merck & Co, Inc.). The vaginal pH was scored by two independent researchers following the manufacturer’s instructions using a scale ranging from 5.5 to 9.0.

Immobilized male rhesus monkeys were placed in ventral recumbency and sampled for urethral swabs. A minitip FLOQ swabs (Copan Improve Diagnostics) was moistened using sterile physiological saline solution and subsequently inserted 1-2 cm into the urethra of the animal. Subsequent handling of the samples was identical to the procedure described for vaginal swab samples.

Suitable precautions were taken during sample collection to avoid microbial contamination. As a sample collection control, a FLOQ swab with a single drop of sterile physiological saline solution was immediately transferred into a 500 µl custom-made lysis buffer at the breeding facility at the time of sampling.

### DNA Extraction

We used the QIAamp Mini Kit (Qiagen GmbH) to extract bacterial DNA. This kit was previously validated for microbial analysis of swab material (*30*). Briefly, proteinase K (50mg/μl) was added and the samples were incubated overnight at 56°C at 600 rpm (Thermomix comfort, Eppendorf). Appropriate amounts of AL buffer (Qiagen GmbH) and ethanol were added. The DNA was subsequently purified from the lysate using the spin columns provided in the kit. Extracted DNA was eluted in 75μl Microbial DNA-Free water (Qiagen GmbH). Suitable precautions were taken during sample handling and processing in the laboratory to limit microbial contamination and maintain consistency during all procedures. The order of sample processing was randomized to avoid handling bias. As a laboratory analysis collection control, a FLOQ swab was transferred into a 500 µl custom-made lysis buffer under the DNA extraction bench at the time the rhesus monkey samples were handled.

### 16S ribosomal RNA gene sequencing

The universal primers 515F and 806R, which were adapted with linker regions and barcode sequences, were used to amplify the V4 region of the 16S ribosomal RNA (16S rRNA) gene (*32*). Phusion Hot Start II High-Fidelity DNA Polymerase (Thermo Fisher Scientific), which has been previously validated for the use in microbiome studies (*30*), was used to amplify each sample in triplets. PCR reactions consisted of 12.5μl of 2x PCR master mix, 8μl of Microbial DNA-Free water (Qiagen GmbH), 1.25μl of each primer (0.5mM each, Metabion) and 2μl of template in a total reaction volume of 25μl. PCR cycling conditions comprised of a pre-denaturation step of 30s at 98°C, followed by 30 cycles of 98°C for 10s, 55°C for 15s and 72°C for 60s, as well as a final 10 min extension step at 72°C. A blank control (Microbial DNA-Free water) and a mock control sample (Microbial mock community, HM-280, Biodefense and Emerging Infectious Research (BEI) Resources, Manassas, USA) were included in 16S rRNA gene amplification. The amplicon triplets were pooled, purified using 0.7x AMPure XP beads (Beckman Coulter), and quantified using the Qubit 2.0 Fluorometer (Thermo Fisher Scientific). Subsequently, we verified the amplicon integrity for a representative number of eleven samples using the BioAnalyzer 2000 (Agilent). Equimolar amounts (10nM) of sample amplicon and maximum volume of control samples (5μl) were pooled prior to sequencing. Illumina MiSeq 2×250bp paired-end sequencing (Illumina V2 chemistry) was performed in the Transcriptome and Genome Analysis Laboratory at the University of Göttingen in accordance with published guidelines (*32*). All generated read files are available at the NCBI Sequence Read Archive (PRJNA521516).

### Data processing and analysis

The sequencing reads were processed using the mothur software package (v.1.39.5) (*33*). According to the MiSeq SOP (*33*), contigs were assembled, sequences were quality filtered, and PCR artifacts were removed. The SILVA bacterial reference database (*34*) was used to align the sequences and OTUs were assigned based on 97% sequence similarity. Cross-sample singletons and poorly aligned sequences were removed. The seq.error command was used to determine the error rate and the mock community was eliminated from the dataset. Due to low read numbers, control sample reads were excluded from the dataset and analyzed separately.

To examine differences in the microbial community structure, alpha (species richness within a single sample) and beta diversity (microbial community diversity between samples) was calculated. As alpha diversity measurements, we determined the number of observed OTUs and calculated the inverse Simpson Metrix using the summary.single command in mothur. Beta diversity was determined using unweighted UniFrac metrics (*35*). The dissimilarity matrix was visualized using Principal Coordinates Analysis (PCoA) and a Newick formatted dendrogram (visualized in FigTree v.1.4.2, http://tree.bio.ed.ac.uk/software/figtree/). ClustVis tool (https://biit.cs.ut.ee/clustvis/) was used to create a heatmap of the relative abundance of bacterial taxa (*36*). Violin plots (R package plot.ly) and box plots (GraphPad Prism 6) were used to visualize data points for different variables.

### Statistical analysis

The statistical significance of the pooled data was analyzed in GraphPad Prism 6 (GraphPad software) and the R package ‘vegan’. Whenever appropriate, we tested for normality distribution of the data using the Kolmogorov-Smirnov normality test. The significance in alpha diversity and pair-wise beta diversity between two or more groups was tested using the non-parametric Mann-Whitney-U or Kruskal-Wallis tests including correction for multiple testing using Dunn’s post hoc tests. Differences in community structure based on age of animals, group association, lactation status and dominance rank was tested using analysis of molecular variance (AMOVA, 1,000 permutations) in mothur (*37*). Principal Coordinates Analysis (PCoA) plots of unweighted UniFrac metrics and UPGMA-clustered dendrograms (unweighted UniFrac metrics) were used to visualize data points. Differences in the ten most abundant OTUs in vaginal samples were assessed using the metastats command in mothur (*38*). p-values for differences in individual OTUs were corrected for multiple comparisons using Bonferroni correction. Values of p < 0.05 were considered statistically significant.

## Acknowledgments

The authors thank the Biodefense and Emerging Infectious Research (BEI) Resources, NIAID, NIH for providing the cells from Microbial Mock Community (Even, HM-280) as part of the Human Microbiome Project. We thank Tamara Becker, Annette Schrod, Uwe Schönmann, Annette Husung, Melina Urh, and all the animal caretakers for their assistance during sample collection. Additionally, we thank Uwe Schönmann and Dietmar Zinner (German Primate Center) for their guidance and general support. We thank the staff of the Transcriptome and Genome Analysis Laboratory at the University of Göttingen for their assistance in optimizing the sequencing run.

## Author contributions

L.H.W., C.R., and S.K designed the study. Sample collection was performed by L.H.W., S.L. and S.K. Laboratory work was conducted at the German Primate Center and performed by L.H.W. and S.L. Data were analyzed by L.H.W. and S.K. All authors (L.H.W., S.L., C.R. and S.K.) contributed to the manuscript preparation.

## Conflicts of Interest

The authors declare no competing interests.

## Data and materials availability

All data will be released after peer-review publication of this article.

